# Macrophages Promote Tumor Cell Extravasation across an Endothelial Barrier through Thin Membranous Connections

**DOI:** 10.1101/2023.02.16.528161

**Authors:** Alessandro Genna, Camille L. Duran, David Entenberg, John Condeelis, Dianne Cox

## Abstract

Macrophages are important players involved in the progression of breast cancer, including in seeding the metastatic niche. However, the mechanism by which macrophages in the lung parenchyma interact with tumor cells in the vasculature to promote tumor cell extravasation at metastatic sites is not clear. To mimic macrophage-driven tumor cell extravasation, we used an *in vitro* assay (eTEM) in which an endothelial monolayer and a matrigel-coated filter separated tumor cells and macrophages from each other. The presence of macrophages promoted tumor cell extravasation while macrophage conditioned media was insufficient to stimulate tumor cell extravasation *in vitro*. This finding is consistent with a requirement for direct contact between macrophages and tumor cells. We observed the presence of Thin Membranous Connections (TMCs) resembling similar structures formed between macrophages and tumor cells called tunneling nanotubes which we previously demonstrated to be important in tumor cell invasion *in vitro* and *in vivo* (Hanna 2019). To determine if TMCs are important for tumor cell extravasation, we used macrophages with reduced levels of endogenous M-Sec (TNFAIP2), which causes a defect in tunneling nanotube formation. As predicted, these macrophages showed reduced macrophage-tumor cell TMCs. In both, human and murine breast cancer cell lines, there was also a concomitant reduction in tumor cell extravasation *in vitro* when co-cultured with M-Sec deficient macrophages compared to control macrophages. We also detected TMCs formed between macrophages and tumor cells through the endothelial layer in the eTEM assay. Furthermore, tumor cells were more frequently found in pores under the endothelium that contain macrophage protrusions. To determine the role of macrophage-tumor cell TMCs *in vivo*, we generated an M-Sec deficient mouse. Using an *in vivo* model of experimental metastasis, we detected a significant reduction in the number of metastatic lesions in M-Sec deficient mice compared to wild type mice. There was no difference in the size of the metastases, consistent with a defect specific to tumor cell extravasation and not metastatic outgrowth. Additionally, examination of time-lapse intravital-imaging (IVI) data sets of breast cancer cell extravasation in the lung, we could detect the presence of TMCs between extravascular macrophages and vascular tumor cells. Overall, our data indicate that macrophage TMCs play an important role in promoting the extravasation of circulating tumor cells in the lung.

## Introduction

During the past several decades much effort and focus has been put into fighting primary breast cancer growth, which has led to an increase in survival rate (90% five-year relative survival) for patients with non-disseminated tumors. In contrast, patients suffering from metastatic disease have a very low (28% five-year) relative survival rate (1). The poor outcome in cases of metastatic disease has led to an intense study of the metastatic process. Metastasis results from a sequence of events that starts in the primary tumor and allows tumor cells to intravasate into the blood stream and use the circulatory system to reach distant organs (2-4). Viable circulating tumor cells (CTCs) that extravasate into the parenchyma of distant organs can then either colonize the organ (leading to overt metastases) or survive in a dormant state while still retaining the ability to produce metastases at a later time, depending on the microenvironment (5, 6).

Macrophages are an important player involved in the metastatic process. Macrophages are very diverse and contain many subpopulations that fulfill different functions including tissue repair, mediation of inflammatory responses, and promotion of tumor progression. Many studies show a strong correlation between high infiltrating macrophage levels in the primary tumor and increased metastatic burden (7, 8). Macrophages aid tumor cells in the primary tumor by promoting tumor cell growth, migration, and invasion, and are referred as tumor associated macrophages (TAMs). TAMs have been shown to secrete EGF and co-migrate in streams with tumor cells towards blood vessels (9-11). In the perivascular region, a subpopulation of Tie2^high^ expressing macrophages, along with stationary tumor cells in a structure identified as a TMEM doorway, play an important role in the intravasation of tumor cells. These macrophages mediate intravasation by creating a local transient opening in the blood vessel (12). Perivascular macrophages are also responsible for inducing tumor stemness and dormancy that prepares the tumor cells to colonize and survive at distant organs (13) (5). Other data indicating the importance of macrophages in metastases is the requirement of colony-stimulating factor 1 (CSF-1) for metastatic dissemination, an essential factor for macrophage survival and proliferation (14, 15). Furthermore, a unique population of macrophages called Metastases Associated Macrophages (MAMs) characterized by the surface markers F4/80+/CSF-1R+/CD11b+/Gr1-CX3CR1^high^/CCR2^high^/VEGFR1^high^ are recruited to the lung and are important for the number and size of metastases in the PyMT experimental metastasis model (16, 17). Furthermore, MAMs are required for efficient metastatic seeding, and present a specific expression pattern that promotes their accumulation and metastatic seeding (18, 19).

It was previously noted in time-lapse intravital images, that tumor cell-macrophage interactions dramatically increased during tumor cell extravasation (20). This interaction occurred despite the fact that 70% of the tumor cells were still located in the vasculature. Additionally, when tumor cells were captured in the act of extravasation (crossing the vessel) macrophages appeared to directly interact with the extravasated part of the tumor cell (21, 22). Additionally, when in association with tumor cells, macrophages extended extremely long, thin pseudopods that would not be visible at lower resolution (21).

The existence of membranous connections between cells *in vivo* was first demonstrated by Chinnery in 2008, describing membranous nanotubes between immune cells and stromal cells in the mouse cornea (23). Similar structures can be found connecting perivascular macrophages to resident tissue cells and between pericytes on separate capillary systems (24, 25). Membranous connections have been described in various tumor types (26). In glioblastoma, they play a role in chemoresistence, and similar structures have been found by light sheet microscopy to be present in MDA-MB-231 brain metastases (27, 28). These structures are often called by different names such as membrane nanotubes (23), tumor microtubes (27), or tunneling nanotubes (TNTs) (29, 30) and can be distinguished by diameter, cytoskeletal components, and whether they are closed-ended or open-ended. However, they all fall under the broad category of thin membranous connections (TMCs).

These structures have been proven to have the ability to promote different functions such as exchanging mitochondrial and nuclear material, potentiating growth factor signaling, and promoting ion exchange (28, 31-33). All these different functions are mediated by establishing direct cell-to-cell contact and communication between distant cells (34). M-Sec (TNFAIP2) is an important regulator of TNT-like membranous connections in macrophages, tumor cells, and other cell types (35). M-Sec is a 73 KDa cytosolic protein whose N-Terminal polybasic region is responsible for M-Sec recruitment to the plasma membrane and its C-terminus recruits RalA, thereby regulating the exocyst function needed for TMC formation (36, 37). Our previous work has shown that macrophage TNTs enhance tumor cell invasion *in vitro* and in a zebrafish model of invasion (33).

Despite the importance of macrophages in tumor cell extravasation, the events leading to extravasation have not been completely elucidated. Here our results using an in vitro assay that mimics extravasation, demonstrate that macrophage TMCs play a major role in promoting tumor cell extravasation. We used an M-Sec deficient mouse (whose macrophages have a defect in TMC formation) to confirm the importance of macrophage TMCs *in vivo*. Furthermore, using high-resolution intravital-imaging (IVI) to examine the interaction of macrophages with tumor cells during extravasation, we found that macrophages in the lung parenchyma can interact with tumor cells via a TMC prior to extravasation.

## Materials and Methods

### Cell Lines and Culture media

MDA-MB-231 cells (ATCC) were cultured in DMEM supplemented with 10% FBS and antibiotics. Where mentioned, MDA-MB-231 cells were serum-starved in DMEM supplemented with 0.5% FBS/0.8% BSA 16□h prior to the start of the experiment. Human umbilical vein endothelial cells (HUVECs, Lonza, Allendale, NJ, USA) were cultured in EGM-2 (Lonza, Allendale, NJ, USA) and only used between passage 1–4. All cells were cultured and maintained in a 5% CO_2_ and 37°C incubator. Murine monocyte/macrophage RAW264.7 subline LR5 (38) was cultured in RPMI 1640 medium (Mediatech) supplemented with 10% heat-inactivated newborn calf serum (Sigma-Adrich) and antibiotics. Bone marrow-derived macrophages (BMMs) were isolated from the tibia and femors of genotyped female mice as described (39) and grown in αMEM with 15% heat-inactivate FBS, 36 ng/ml recombinant human CSF-1 (kindly provided by ER Stanley), and antibiotics. E0771-GFP medullary breast adenocarcinoma cells, originally isolated from a spontaneous mammary tumor in C57BL/6 mice, were obtained from Dr. Wakefield’s lab at the NIH, who in turn obtained them from Dr. Fengzhi Li in Dr. Enrico Mihich’s lab at Roswell Park Cancer Institute, Buffalo, NY.

### Antibodies and reagents

Most reagents including Cell-Tracker Red (CMPTX) cat # C34552 and Cell-Tracker Green (CMFDA) cat# C7025 were obtained from Thermo Fisher. ZO-1 Polyclonal Antibody (Invitrogen, cat#40-2200) and M-Sec /TNFAIP2 (AbCam, ab91235) antibodies were used for immunofluorescence and anti-actin (Sigma) or anti-TNFAIP2 (Invitrogen, PA5-13542) were used in western blotting.

### Mice

All procedures were conducted in accordance with the National Institutes of Health regulation concerning the care and use of experimental animals and with the approval of Albert Einstein College of Medicine Animal Care and Use Committee. The TNFAIP2 KO mouse model was generated by CRISPR technology. In brief, a guide RNA (gRNA) targeting exon2 of TNFAIP2 gene, Tnfaip2 Ex2 gRNA 62/85 (targeting sequence: aaagccacgctctacatccgagg) was designed by online software Benchling (www.benchline.com) and generated by in vitro transcription. Cas9 protein was purchased from PNB. All the above CRISPR ingredients were injected into the fertilized eggs of C57BL6 mice and then the injected fertilized eggs were transferred to the pseudopregnant CD1 female mice for production of offspring. TNFAIP2 genotyping was done using primers F: AGCTGTGCATTTAGGTCCAT and RW: TGTGGGCAGTGGACCATCTA. Loss of M-Sec protein was confirmed by western blotting using an M-Sec specific antibody. Homozygous M-Sec–KO mice were viable, born at normal Mendelian ratios, grew normally, and showed a normal phenotype similar to that reported using a M-Sec deficient mouse generated by the Riken Centre (40). For the intravital imaging C57Bl6 mice were generated by crossing Tg(Csf1r*-GAL4/VP16,UAS-ECFP)1Hume/J (Stock No: 026051, Jackson Laboratory) mice with B6.Cg-Gt(ROSA)26Sortm14(CAG-tdTomato) to label the vasculature with tdTomato VE-cadherin, as described in (22).

### Western Blot

Cells were lysed in ice-cold buffer containing 25 mM Tris-HCl, 137 mM NaCl, 1% Triton X-100, 2 mM EDTA, 1 mM orthovanadate, 1 mM benzamidine, 10 *μ*g/ml aprotinin, 10 *μ*g/ml leupeptin, and pH 7.4. Whole-cell lysates were resolved by SDS-PAGE, and proteins were transferred onto PVDF membranes (Immobilon-P, Millipore) that were subsequently blocked using 3% BSA, 1% ovalbumin in TBS containing 0.1% Tween20 before incubation with primary antibodies overnight at 4°C. Membranes were then washed and incubated with secondary antibodies conjugated to HRP. Images were acquired using a Kodak Image Station 440.

### Extravasation Trans-Endothelial Migration (eTEM) assay

8-micron pore sized transwell inserts (Millipore) were coated with 50μL of 300 μg/mL of growth factor reduced matrigel (BD) diluted in DMEM media (Invitrogen) for 1 hour at 37°C. Excess matrigel was removed and 100K HUVECs were plated on the matrigel-coated filters in 300μL EGM-2. Transwells were incubated in a 24 well plate containing media at 5%CO_2_/37°C for 48h. Cell tracker-red labeled macrophages were added to the underside of the transwell containing an endothelial monolayer and allowed to adhere for 2 hours at 5%CO_2_/37°C. Transwells were then returned to the 24 well plate and serum starved cell tracker green-labeled tumor cells were added into the upper side of the chamber and incubated at 5%CO_2_/37°C. To assess the extent of tumor cell trans-endothelial migration, after 18h transwells were washed twice with PBS and then fixed in 4% paraformaldehyde/PBS for 20 minutes, permeabilized 0.1% triton X-100 for 10 minutes and stained with ZO-1 (Invitrogen). Transwells were placed in PBS on a Mattek dish (MatTek Corp) and imaged on a previously described custom built inverted multiphoton microscope with a 25X NA 1.05 water immersion objective and a Spectra Physics Mai Tai-DeepSee laser set to 880nm (41). 1μm step Z-stacks views were acquired spanning the depth of the transwell, and only those tumor cells that breached the endothelium were scored as trans-endothelial migration events.

### TMP quantitation and proximity analysis

The ability of either macrophages or tumor cells to make a thin membranous protrusion (TMP) across the filter and the endothelium was determined and quantified. In brief, the eTEM assay was assembled as described above and fixed at 1h, 2h, and 3h after tumor cell seeding. The transwells were than permeabilized, stained and imaged. In order to be scored as a TMP in the eTEM assay, the cell protrusion has to cross the filter while the main cell body remains on the originally seeded face of the chamber. To determine the probability of finding tumor cells in proximity of macrophages, a circle with a radius of 30 µm was drawn around a pore with macrophages on the extra-vasculature space, which was defined as a region of interest (ROI). The ROI was then imported into the corresponding image of the vascular side of the transwell and the presence of tumor cells within the circle was scored.

### TMC quantitation

The ability of cells to make TMCs *in vitro* was monitored following culture of RAW/LR5 or BMMs overnight in MatTek dishes (MatTek Corporation) in complete growth medium. Cells were briefly washed with PBS buffer and then imaged live so as to avoid the TMC breakage that occurs during fixation. In order to be counted as a TMC connection, at least one TMC of at least 8 µm in length was required to be present between two cells with at least a portion of the TMC not adherent to the substratum. To be categorized as TMC negative, cells without TMCs were required to be within one cell body length of another cell without touching any other cell. TMCs were also quantified in the eTEM assay. The eTEM assay was performed as described above (see in vitro extravasation assay) and fixed after 2 hours.

### Quantification of Lung Metastasis

E0771-GFP tumor cells were prepared by trypsinizing a 10□cm confluent culture dish and passing the cells through a 40 μm cell strainer (Falcon, cat #352340) to remove clumps. A total of 2□×□10^5^ cells was resuspended in 100□μL of sterile PBS and intravenously (iv) injected via the lateral tail vein. Lungs were harvested 1 week after injection and GFP positive cells were counted by epifluorescence microscopy using a 20X phase objective on an Olympus IX71 microscope. The number of metastatic foci of >5 cells was scored. Images of nodules were taken using a Sensicam cooled CCD camera mounted on the microscope. Images were thresholded in ImageJ (National Institutes of Health) and the size of nodules was quantified using the analyze particles function.

### Window for high resolution imaging of the lung (WHRIL) and intravital imaging

The surgery for the WHRIL implantation and the Window passivation method were performed as described previously (22, 42). Briefly, mice are anesthetized, the skin and muscle resected from the upper left chest region and a 5□mm circular defect is cut in the chest wall exposing the lung tissue. A window frame is sutured in place and the underside adhered to lung tissue while applying positive end-expiratory pressure. Next a 5 mm coverslip is coated on one side with a thin layer of adhesive and used to seal the aperture of the window frame and attach the exposed lung tissue to the glass. A purse-string suture is used to cinch the ribs within the window frame groove and another purse-string suture is used to cinch the skin within the same groove. Finally, an insulin syringe is placed into the thoracic cavity through the diaphragm and used to remove any excess air. After induction of anesthesia using 5% isofluorane, windows were captured within a fixturing plate, animals were placed on the microscope stage and the fixturing plates were taped to the stage using paper tape. The animal was placed in a heated chamber maintained at physiological temperatures by a forced-air heater (WPI Inc., AirTherm ATX), during the course of imaging and the animals were maintained at 0.75-1.5% isoflurane for the duration of imaging. Imaging was performed on a previously described, custom-built, inverted multiphoton microscope equipped with a 25X NA 1.05 water immersion objective and a Spectra Physics Mai Tai-DeepSee laser set to 880nm (41). Z-stacks (21 µm deep, 3 µm step size) time lapse imaging (one stack every 1.4 min for 8 h) was acquired using two frame averages in 16□bit using a 25×1.05 NA objective lens.

## Results

### Macrophage TMCs are important in promoting tumor cell extravasation *in vitro*

Macrophages play an important role in promoting metastatic seeding (18, 19) and we previously detected stromal macrophages in contact with tumor cells arriving at the metastatic site (22). However, it is unclear how macrophages in the lung parenchyma interact with CTCs in the vasculature. To specifically study the role of TMCs connecting macrophages to tumor cells through an endothelium, we employed an *in vitro* assay that mimics macrophage assisted tumor cell extravasation, maintaining the physical characteristics, the physiology, and cellular orientation of macrophages and tumor cells in the lung vasculature (Fig. 1A)(16). We cultured a highly metastatic triple negative breast cancer cell line (MDA-MB-231) on a transwell membrane coated with an intact layer of endothelial cells. The endothelial layer separated the tumor cells from cells of a murine derived macrophage cell line (RAW 264.7/LR5) that were attached to the bottom of the filter. We could detect a small but insignificant amount of tumor cell crossing into the extravascular space (bottom of the chamber) starting an hour after the addition of tumor cells (Fig. 1B). During this period we observed macrophage membrane extensions spanning the endothelial layer and the filter’s pores connecting to tumor cells (TMCs). TMCs could be detected two hours post tumor cell seeding (Fig. 1C-D). We then determined the frequency of tumor cells or macrophages to make thin membranous protrusions (TMPs) or connections (TMCs) into the layer between the two cell types over time. While not significant, macrophages seem to generate a greater number of protrusions as compared to tumor cells (Fig. 2A). We also investigated the conditions under which these macrophage protrusions occur in response to tumor cells. We determined whether there was an association between macrophages forming protrusions and the presence of tumor cells on the upper side of the endothelium in the vascular space. We measured the association of macrophages with or without protrusions through a pore towards the endothelial layer with the presence of tumor cells on the vascular side of endothelium within a diameter of 60 µm centered on the pore (Fig. 2B). Our data shows that tumor cells are found more frequently associated to areas with macrophages containing protrusions (86%) while, there was only a random association between macrophages without protrusions and tumor cells (48%) (Fig. 2C-D). These data indicate that tumor cells are preferentially localized to areas above macrophages containing protrusions. However, it is unclear whether macrophage protrusions are being directed towards tumor cells, or whether tumor cells migrate towards sites above macrophage with protrusions.

**Figure 1:**
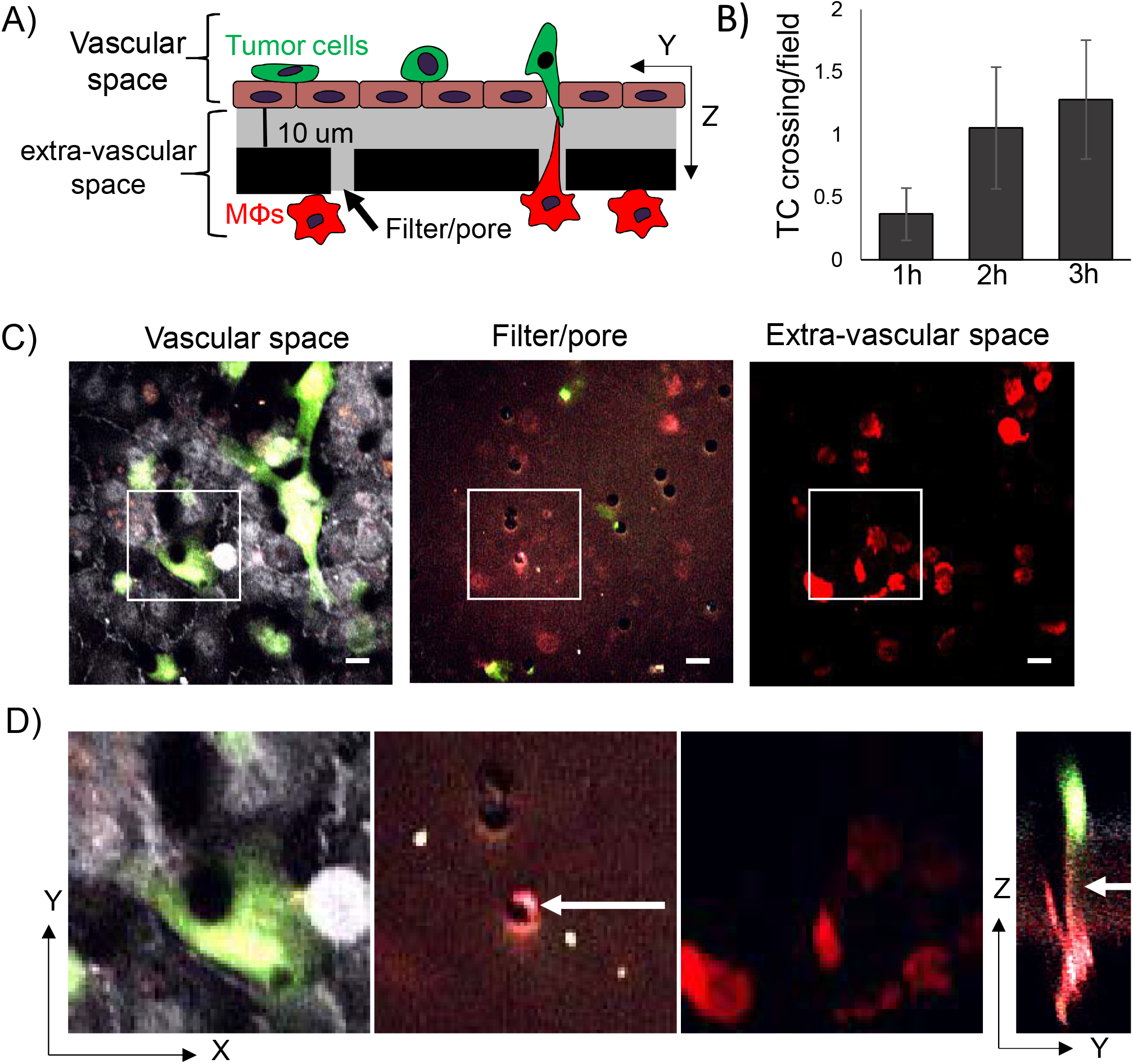
Macrophage TMCs connecting to tumor cells can be detected using an in vitro tumor cell extravasation transendothelial migration (eTEM) assay. A) Schematic representation of eTEM assay. The vascular space is contained in the upper part of the chamber and contains green labeled tumor cells on top of a confluent endothelial layer. The endothelial layer is supported by a matrigel (grey) coated filter with 8 µm pores with red labeled macrophages (MΦs) on the underside of the filter (the extra-vascular space). B) The number of tumor cells (TC) crossing the endothelium over time was determined as the number per field in the extravascular space. C) Representative images of the vascular space, the supportive filter, and the extravascular space. Tumor cells and macrophages are labeled using cell tracker green and cell tracker red, respectively, and endothelial cells are stained for the tight junction marker ZO-1 (white). Scale bar 15 μm D) Magnified images of the boxed areas in C) and an orthogonal (Y-Z) view of the Z-stack marked by boxed area in figure C.

**Figure 2:**
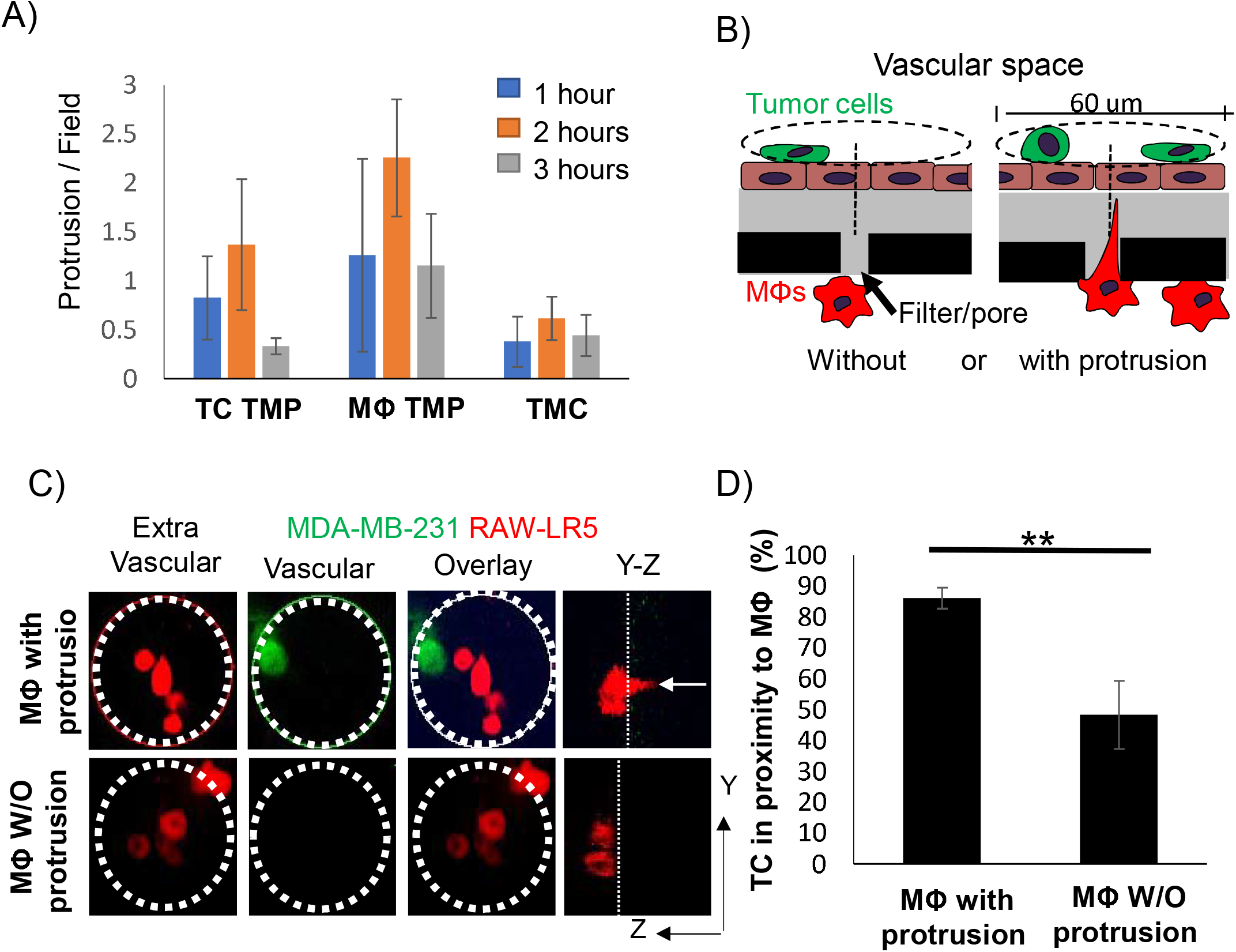
Tumor cell localization to macrophages occurs in high frequency when macrophages contain thin membranous protrusions towards the upper surface. A) Time course of the formation of thin membranous protrusions (TMP) and thin membranous connections (TMC) over time in the eTEM assay. N=3 experiments. B) Schematic of quantification strategy with green tumor cells located on top of the endothelial layer (brown) in the vascular space and macrophages (red) in the extra vascular space. A 60µm diameter circle centered around a pore containing macrophages, either with or without protrusions, and the presence of tumor cells on the vascular surface within the circle (dashed line) was scored. C) Representative images of the macrophages (red) in the extravascular space and tumor cells (green) in the vascular space in the eTEM assay 2 hours after tumor cell introduction, along with an overlay of the two images to determine the co-localization of both cell types. Dotted white circle represents the analyzed area. White arrow in Y-Z orthogonal view marks the macrophage protrusion crossing the filter. D) % of the time that tumor cells are found in proximity (within 30µm radius) to macrophages with and without protrusions. N=3 experiments, ** P value<0.01

The TMCs we detected in the early stage of the extravasation process *in vitro* resembled TNT-like structures that we previously identified to exist between macrophages and tumor cells that were important for tumor cell invasion (33). Therefore, we hypothesized that the known TNT regulator, M-Sec, would affect the ability of macrophages to promote tumor cell extravasation. To test this hypothesis, we used previously generated macrophage cell lines expressing either a control or an M-Sec targeting shRNA (shCtrl and shM-Sec) (43). We extended the time of incubation of our *in vitro* eTEM assay to allow for maximal tumor cell crossing into the extravascular space during extravasation (Fig. 3A-B). We verified the level of M-Sec in shCtrl and shM-Sec macrophages and the defect in TMC formation. There was a significant 50% reduction of M-Sec protein level in shM-Sec macrophages, which corresponded to a 50% reduction in the capability of these cells to make TMCs (Fig. 3C). We then determined the capacity of the shM-Sec macrophages to promote tumor cell extravasation in the eTEM assay, as shown by the analysis of the number of tumor cells crossing the endothelium (Fig. 3D). Tumor cells alone have a baseline ability to extravasate that was significantly increased by the presence of shCtrl macrophages. Although shM-Sec macrophages still promoted tumor cell extravasation, this was significantly reduced compared to control macrophages, which was similar to the level of reduction of M-Sec protein levels in these cells. This result shows that macrophage TMCs play an important role in promoting tumor cell extravasation. Next, we determined whether tumor cell extravasation was dependent on soluble factors secreted by macrophages. Macrophage conditioned media was unable to promote a significant increase in tumor cell extravasation above the baseline level of tumor cells alone (Fig. 3D). Overall, these data indicate that macrophages contact tumor cells through TMCs and reduction in the production of TMCs reduces tumor cell extravasation *in vitro*.

**Figure 3:**
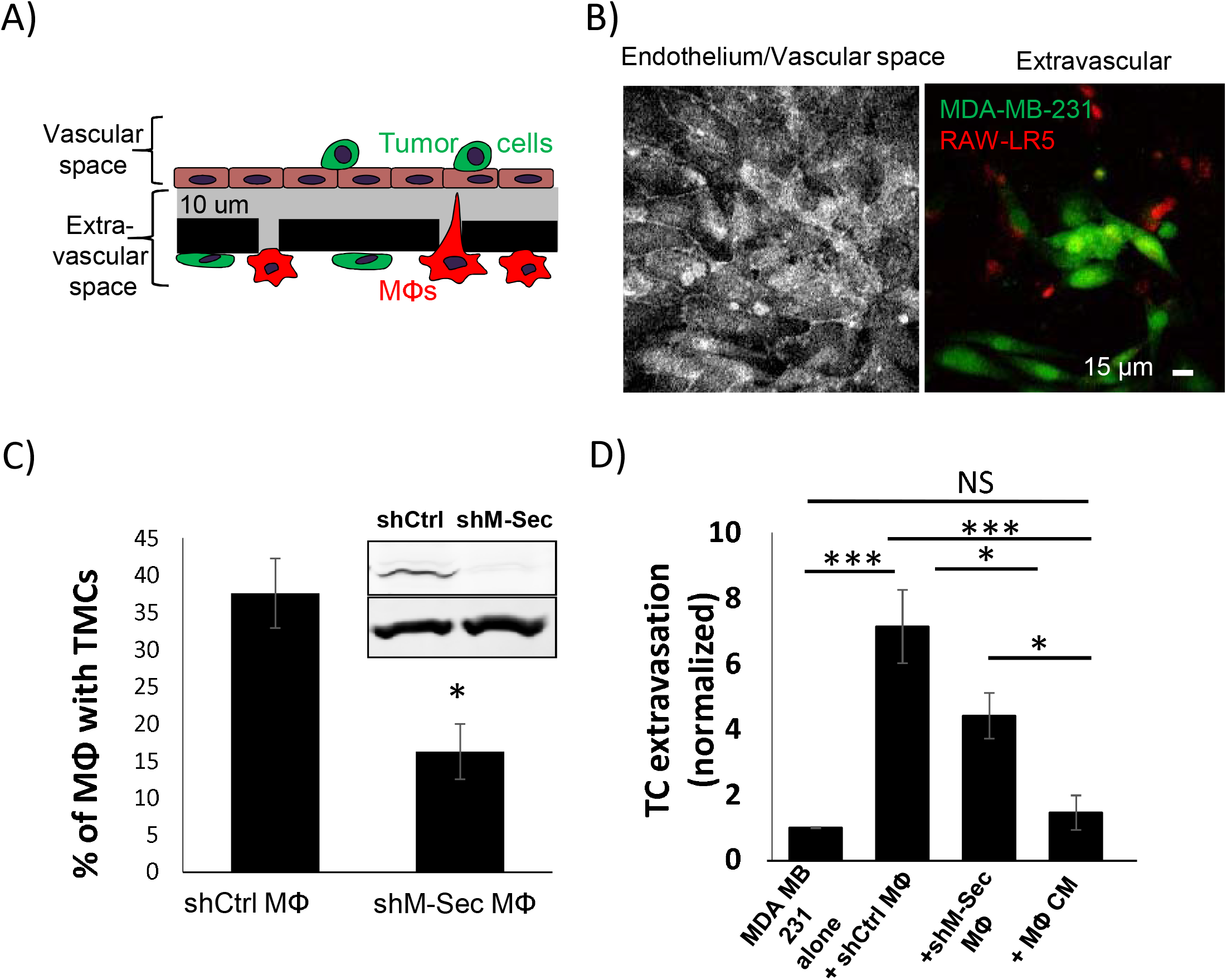
Macrophages defective in TMCs do not promote tumor cell extravasation. A) Schematic representation of eTEM assay after overnight incubation to allow for maximal tumor cell crossing. The vascular space is contained in the upper part of the chamber and contains green labeled tumor cells on top of a confluent endothelial layer. The endothelial layer is supported by a matrigel (grey) coated filter with 8 µm pores with red labeled macrophages (MΦs) on the underside of the filter (the extra-vascular space. B) Representative images of the intact endothelium stained for ZO-1 (left) in the vascular space and (right) the tumor cells (green) that have crossed into the extravascular space 24 hours after tumor cell addition. C) shRNA Control (shCtrl MΦ) or shRNA M-Sec (shM-Sec MΦ) macrophages were cultured overnight and the percentage of macrophage making TMCs between cells quantified. Inset - Western Blot showing M-Sec protein (upper) and actin (lower) level in macrophages expressing either shRNA control (shCtrl) or shRNA M-Sec (shM-Sec). D) The number of MDA-MB-231 tumor cells (TC) crossing the endothelial layer into the extravascular space after 24 hours in absence (alone) or presence of RAW/LR5 macrophages that express shRNA Control (shCtrl MΦ) or shRNA M-Sec (shM-Sec MΦ), or in the presence of MΦ conditioned media (CM). The number of TCs present in the extravascular space, normalized to the number of TCs crossing alone. 25 fields per condition were analyzed for each of 3 independent experiments. * P < 0.05, *** < 0.001, NS = non-significant. D)

### TMCs promote tumor cell extravasation *in vivo*

To verify the importance of macrophage TMCs *in vivo* we generated an M-Sec deficient mouse (M-Sec KO) using CRISPR. We verified the loss of M-Sec by Western blot analysis of isolated bone marrow derived macrophages (BMMs) from both control and M-Sec KO mice and (Fig. 4A). We also quantified the ability of the BMMs to produce TMCs *in vitro*, which was reduced almost 60% in the M-Sec KO BMMs (Fig. 4B). We also determined whether the reduction in TMC formation in M-Sec KO BMMs in culture was recapitulated in the eTEM assay. As expected, the number of connections between M-Sec KO macrophages and MDA-MB-231 cells was reduced by 70% (Fig. 4C). The remaining TMCs in the M-Sec KO BMMs may be due to residual level of M-Sec being present or because of alternative mechanisms of TMC formation (29, 34). We further investigated if the loss of M-Sec altered the ability of the macrophages to stimulate tumor cell extravasation *in vitro*. In this case, we used a murine metastatic breast tumor cell line expressing GFP (E0771-GFP). In accordance with our hypothesis that macrophage TMCs are critical for the promotion of tumor cell extravasation and our data using the shM-Sec cell line, loss of M-Sec in BMMs also significantly reduced tumor cell extravasation with no significant difference to baseline levels *in vitro* (Fig. 4D). This data confirmed that M-Sec KO BMMs recapitulated the data using the shM-Sec cell line (Fig. 3D) and suggest that the M-Sec KO mouse model can provide valuable *in vivo* data.

**Figure 4:**
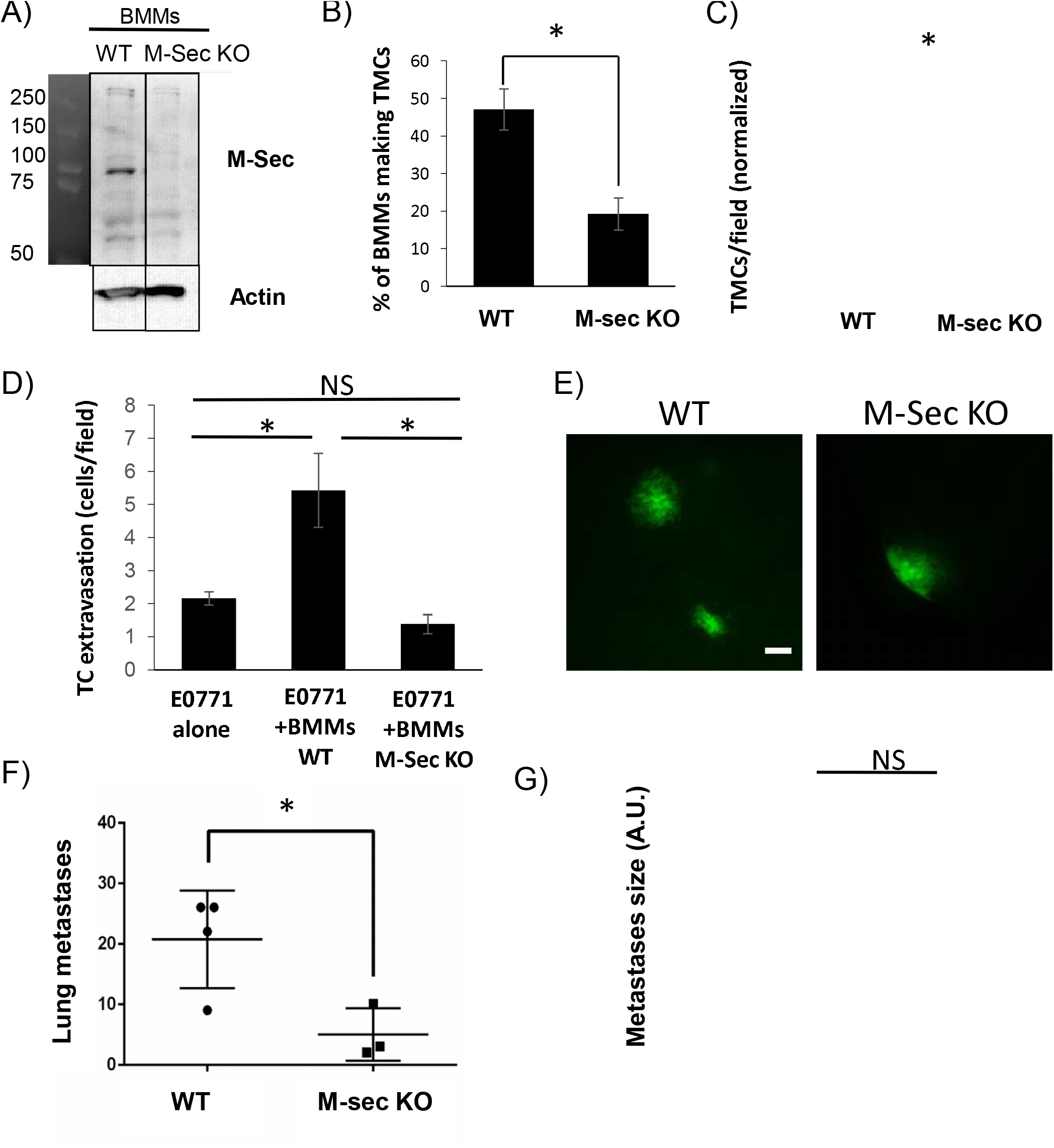
Tumor cell extravasation is facilitated by macrophage TMCs in vitro and in vivo. A) Western Blot showing M-Sec level in bone marrow derived macrophages (BMMs) isolated from either WT or M-Sec KO mice. B) WT or M-Sec KO BMMs were cultured overnight and the percentage of BMMs making TMCs between macrophages and tumor cells determined. C) Quantification of TMCs formed between MDA-MB-231 cells and either WT or M-Sec KO BMMs across the endothelium 2 hours after tumor cell introduction to the eTEM assay (N=3 experiments). D) Quantification of GFP-E0771 tumor cells (TC) crossing the endothelium in the absence or presence of wild type (BMMs WT) or M-Sec deficient (BMMs KO M-Sec) in the eTEM assay. N= 3 experiments, 36 fields per experiment for each condition where analyzed. E) Representative images of lung metastases derived from GFP-E0771 cells injected in the tail vein of either WT or M-Sec KO mice 7 days after tumor cell injection, scale-bar 100 μm. F) Quantification of total number of metastases in either WT or M-Sec KO mice injected with E0771-GFP. G) Quantification of metastasis size, per field. (N=4 for WT and N=3 for M-Sec KO). For all - * P < 0.05, NS = non-significant.

To determine if macrophage M-Sec dependent TMC formation was important for tumor cell extravasation *in vivo*, we injected syngeneic E0771-GFP into the tail vein of either WT or M-Sec KO mice. One week post injection animals were sacrificed, and the lungs analyzed for metastatic burden. The lungs of all mice contained tumor cells on the surface (Fig 4E). Following quantification of metastatic lesions, it was revealed that the M-Sec KO animals had a significantly lower number of metastases compared to the WT animals (Fig 4F). The reduction in metastases was not due to cell growth as there was no significant difference in the size of the individual metastases (Fig 4G).

Previous studies showed that macrophages extend extremely long, thin pseudopods when in proximity to tumor cells (21) but protrusions across the endothelium had not been previously identified. To determine if TMCs occurred during tumor cell extravasation *in vivo* we utilized the WHRIL (window for high-resolution imaging of the lung) with an experimental metastasis model to obtain high resolution images of tumor cell extravasation into the lung (22). E0771-GFP cells were injected into the tail vein and images of the tumor cells in the blood vessels of lungs were collected in 3D over time. We examined z-slices with stationary tumor cells in the blood vessels for the presence of macrophages in the lung parenchyma that extend protrusions through the intact blood vessel endothelium to reach the tumor cell in the blood vessel, creating a TMC (Fig. 5). We observed an interaction between a stationary tumor cell in the vascular space and a macrophage in the extravascular space mediated by a long thin tube which crossed through the endothelium (seen in Fig. 5A) and that this interaction preceded tumor cell extravasation into the lung parenchyma (Fig. 5D). This suggests that macrophages contact tumor cells through TMCs, which may initiate or enhance tumor cell crossing.

**Figure 5:**
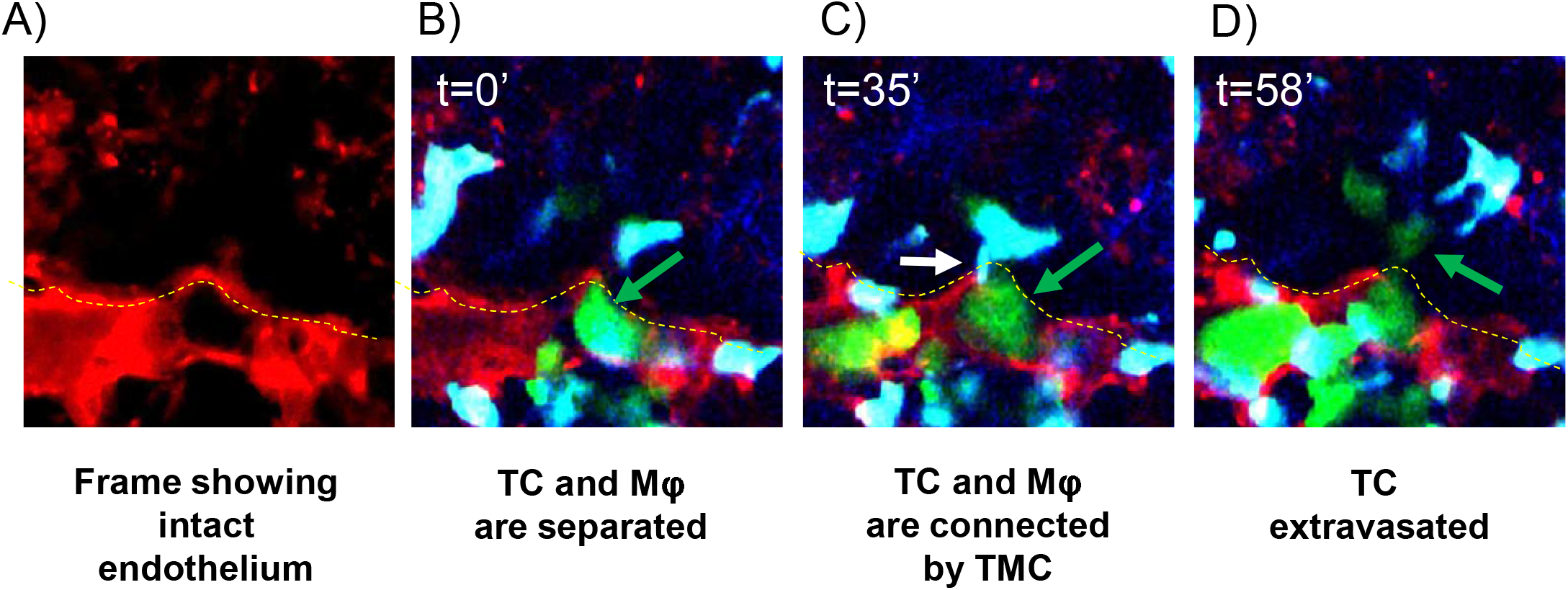
IVI of tumor cell extravasation to the lung reveals the presence of a thin membranous connection (TMC) from a tissue macrophage contacting a circulating tumor cell prior to tumor cell extravasation. A) Representative image showing the intact endothelium (red) of the lung capillary. Images were taken using the WHRIL (window for high-resolution imaging of the lung). B) Representative image of stationary E0771-GFP tumor cell (green) in a lung capillary. Tumor cells were injected through the mouse tail vein in an experimental metastasis model. Yellow dashed line indicates the intact blood-vessel endothelium (red) separating the tumor cell (green) and macrophages (blue) in the parenchyma. C) A TMC (white arrow) protruding from the macrophage and connecting to the stationary intravascular tumor cell in panel B indicated by the green arrow. D) Tumor cell indicated in panels B and C has extravasated into the lung parenchyma (green arrow).

Taken together these data support the importance of TMCs in the metastatic process and, in particular, highlights the possible role of TMCs in promoting the extravasation of disseminated tumor cells.

## Discussion

Metastasis is estimated to be responsible for approximately two thirds of cancer related mortality (44). The importance of macrophages in promoting metastatic dissemination and growth in primary as well as secondary breast cancers is well-known (3, 5, 6). In this work, we focused our attention on the role of macrophages in tumor cell extravasation and, in particular on the physical interaction between macrophages and tumor cells that reside on opposing sides of the lung vasculature. We were able to identify TMCs in an *in vitro* assay that mimics the extravasation process. Similar barrier-crossing membranous extensions have already been described to be made by different immune cells *in vivo*. Dendritic cells send dendrites outside the epithelium to sample bacteria in the intestinal lumen (45) and hematopoietic stem cells send TMCs across the dense basement membranes to deliver lysosomes into diseased proximal tubular cells when grafted to cystinotic kidneys (46). However, we are the first to show a connection between tumor cells and macrophages in the context of the metastases both *in vitro* and their importance in tumor cell extravasation *in vivo*.

Our previous work has shown that similar structures that we identified as TNTs, a subset of TMCs, were important in promoting tumor cell invasion and dissemination (33). We confirmed that the TMCs that we detected during extravasation were regulated in the same manner as TNTs by using macrophages with reduced levels of M-Sec, a known regulator of the TNTs (33, 36, 40, 47). Reduction of M-Sec decreased TMC formation between macrophages and tumor cells and reduced tumor cells extravasation *in vitro*. Taken together these data suggest that physical contact between tumor cells and macrophages promote tumor cell extravasation through TMCs. To further investigate the role of macrophage TMCs in metastatic seeding we generated a mouse model for M-Sec deficiency by introducing a single point mutation in the M-Sec gene through CRISPR technology. BMMs isolated from the M-Sec KO mouse presented a lower capability of making TMCs and were not able to promote extravasation of tumor cells *in vitro*. These results give us confidence that macrophages generated from M-Sec KO mouse recapitulates our results using macrophage cell lines.

Our initial intravital imaging of macrophages interacting with tumor cells during extravasation in vivo suggests that macrophages contact tumor cells through TMCs, which may initiate or enhance transendothelial migration of tumor cells. Additional studies are needed in order to determine the frequency and kinetics of macrophage TMC contact with tumor cells *in vivo* before during and after extravasation.

The importance of M-Sec and TNTs/TMCs *in vivo* was recently demonstrated by another group that showed that M-Sec–TNTs played a protective role in the glomeruli by rescuing podocytes via mitochondrial horizontal transfer (40). We tested the capability of injected wild type tumor cells to disseminate into the lung parenchyma of M-Sec KO mice. The number of metastases in the M-Sec KO mice was significantly reduced compared to wild type mice indicating a role for M-Sec in regulating metastatic dissemination in the lung microenvironment. Taking into consideration that there was no significant change in the size of the metastases we believe that M-Sec generation of TMCs is important for the seeding of circulating tumor cell into the lung parenchyma and not for the growth of seeded tumor cells. Others have shown that factors like Flt-1 regulate the growth (48).

While we showed direct macrophage contact facilitated tumor cell extravasation, which could not be accomplished via secreted factors in conditioned media *in vitro*, this does not mean that there is not a role for secreted factors in the process. For example, MAMs which originate from circulating inflammatory monocytes, are recruited by the CC-chemokine ligand 2 (CCL2) and, once recruited, MAMs secrete another chemokine, CCL3, which leads to MAM retention (19). A number of genes are upregulated in MAMs compared to resident lung or splenic macrophages that may regulate the ability to generate TMCs in macrophages. Ongoing studies are focused on determining the factors regulating macrophage TMCs MAMs. Additionally, tumor cell derived factors may account for the association of macrophage protrusions towards tumor cells on the opposite of the endothelium but defining which chemokine is contributing to this process is a topic for future study.

Even though many aspects of TMCs (such as the molecular mechanisms that drive them or how they promote tumor cells extravasation) are still unknown, our data point towards an important role of TMCs in promoting tumor cell extravasation at the secondary site.

## Acknowledgements

The authors would like to thank the Montefiore Einstein Cancer Center, and the Gruss-Lipper Biophotonics Center and its associated Integrated Imaging Program as well as the Evelyn Gruss-Lipper Charitable Foundation. The authors would like to thank Dr. Lalage Wakefield’s lab at the NCI for the donation of the E0771 cell line.

This work was supported by the NCI grants F32-CA243350 (CLD) and R01-CA216248 (JSC & DE), The Gruss-Lipper Biophotonics Center, The Integrated Imaging Program, The Evelyn Gruss-Lipper Charitable Foundation, and the Integrated Imaging Program for Cancer Research. CLD was partially funded under an K12 GM102779 (DC) IRACDA fellowship where the content is solely the responsibility of the authors and does not necessarily represent the official views of the K12 support.

## Supplemental Figures

**Supplemental Figure 1:**
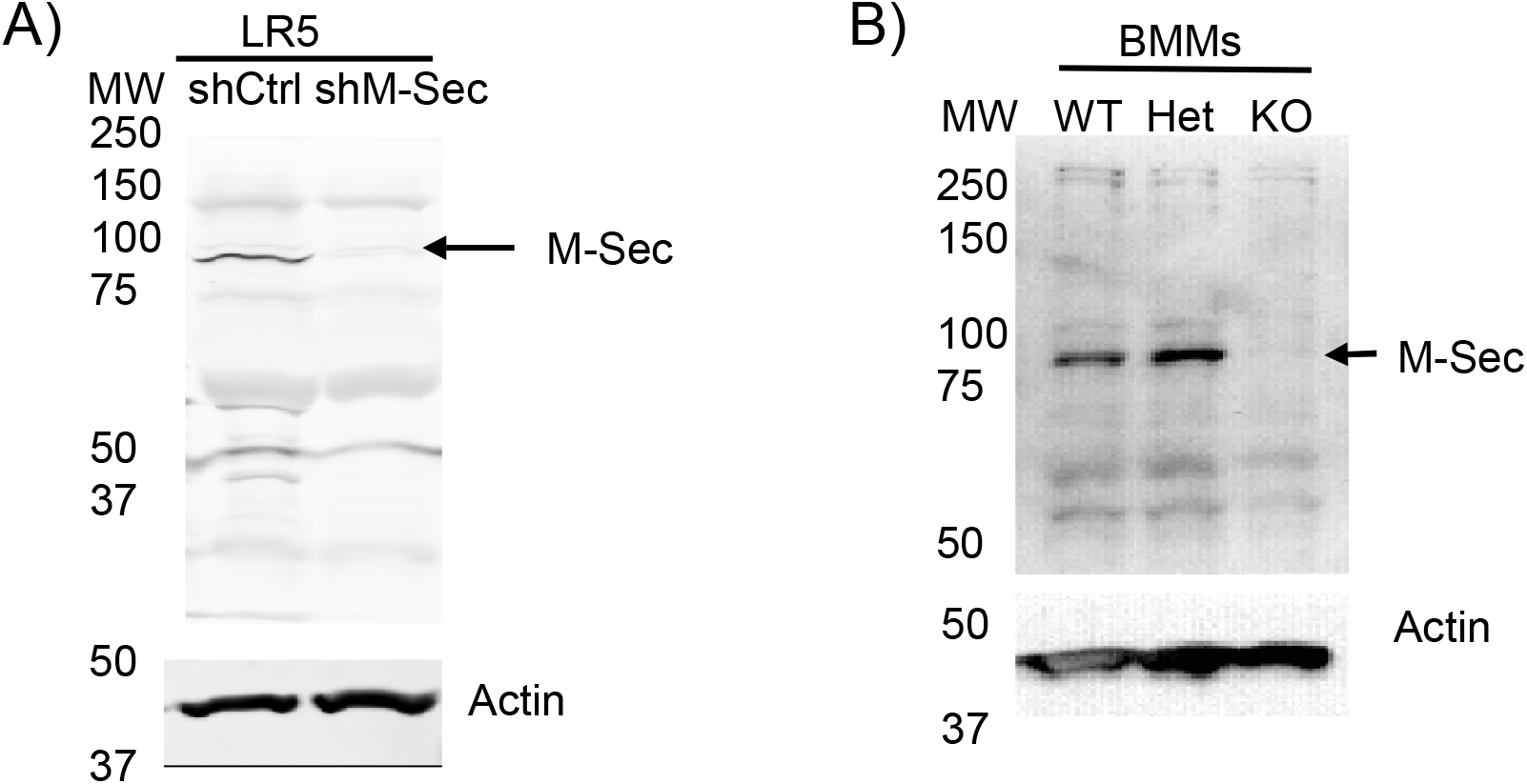
Detection of M-Sec protein levels in the various macrophages used in this study. Western blots of whole cell lysate blotted with specific antibodies against actin (Sigma) for a loading control or M-Sec/TNFAIP2 (Invitrogen, PA5-13542). A) Western Blot showing M-Sec protein (upper) and actin (lower) level in RAW/LR5 macrophages expressing either shRNA control (shCtrl) or shRNA M-Sec (shM-Sec). B) Western Blot showing M-Sec levels in bone marrow derived macrophages (BMMs) isolated from either WT, heterozygous (HET) or M-Sec KO (KO) mice.

## Notes

### Competing Interest Statement

The authors have declared no competing interest.

